# Second Harmonic Generation Spectroscopy of Membrane Probe Dynamics in Gram-Positive Bacteria

**DOI:** 10.1101/645788

**Authors:** L. N. Miller, W. T. Brewer, J. D. Williams, E. M. Fozo, T. R. Calhoun

**Affiliations:** University of Tennessee, Knoxville

## Abstract

Bacterial membranes are complex mixtures with dispersity that is dynamic over scales of both space and time. In order to capture adsorption onto and transport within these mixtures, we conduct simultaneous second harmonic generation (SHG) and two photon fluorescence measurements on two different gram-positive bacterial species as the cells uptake membrane-specific probe molecules. Our results show that SHG can not only monitor the movement of small molecules across membrane leaflets, but is also sensitive to higher-level ordering of the molecules within the membrane. Further, we show that the membranes of *Staphylococcus aureus* remain more dynamic after longer times at room temperature in comparison to *Enterococcus faecalis*. Our findings provide insight into the variability of activities seen between structurally similar molecules in gram-positive bacteria while also demonstrating the power of SHG to examine these dynamics.

**STATEMENT OF SIGNIFICANCE:** Bacterial membranes are highly adept at discerning and modifying their interactions with different small molecules in their environment. Here we show how second harmonic generation (SHG) spectroscopy can track the dynamics of structurally similar membrane probes in two gram-positive bacterial species. Our results reveal behavior that is dependent on both the probe molecule and the membrane composition. Specifically, we observe flip-flop between leaflets for one molecule, while the other molecule produces a signal indicative of larger scale ordering in the membrane. These phenomena can all be explained by considering potential differences in the membrane fluidity and surface charge between the two bacterial species. Overall, our work highlights the dynamic differences between bacterial membranes and SHG’s sensitivity to probing these systems.

## INTRODUCTION

The vast majority of antibacterial drugs are small molecules (1). In order to understand the reasons behind their successes and failures, it is necessary to first understand their initial interaction with bacterial cells, and more specifically, the bacterial membrane which is responsible for regulating small molecule uptake. The membrane of each bacterial species, however, is composed of an unique combination of proteins and phospholipids which can significantly alter its interaction with small molecules. Further, bacterial membranes are dynamic and can adapt their lipid compositions in response to environmental changes (2).

A specific system that highlights the ramifications of variable membrane compositions on drug outcomes is recent work studying the lipopeptide antibiotic, daptomycin. When gram-positive *Staphylococcus aureus* and *Enterococcus faecalis* were grown in media supplemented with oleic acid, *E. faecalis* showed increased tolerance against daptomycin, while *S. aureus* exhibited increased susceptibility (3, 4). The underlying mechanism behind this contrasting activity is currently unclear. Multiple studies have investigated the composition of the membranes of these two bacteria and their differences (3–8), but there is significantly less work in the field examining the broader implications of how these differences impact small molecule interactions. Here, we measure second harmonic generation (SHG) simultaneously with two-photon fluorescence (TPF) to examine differences between these bacterial species *in vivo* as well as their sensitivity to small structural changes to the adsorbing molecules. Through the combination of these methods, we have access to the kinetics of the initial molecule-membrane interaction and subsequent transport processes, thus providing an avenue to describe the underlying mechanisms of action.

Second harmonic generation (SHG) is a nonlinear optical technique that is sensitive to molecular species at interfaces and has previously been been applied to the study of small molecules interacting with both model (9–21) and living cell membranes (22–29). In SHG, incoming light at a frequency, *ω*, with sufficient intensity induces a second order polarization in the sample. This induced polarization stimulates the radiation of new light at twice the incident frequency (2*ω*). Given the second order-field interaction, randomly oriented molecules in centrosymmetric and isotropic bulk media generate SHG signals that, on average, destructively interfere. This leaves molecules that are aligned at interfaces, such as membranes, radiating a constructively interfering SHG signal that can be detected. This interference gives rise to SHG’s exceptional interfacial specificity. The radiated SHG intensity from a single bacterial interface is proportional to the absolute square of the two driving laser fields (*E*_*ω*_) and the effective second order susceptibility, 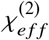:

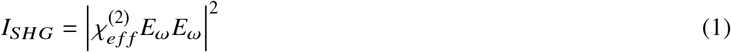

The effective second order susceptibility can further be expressed as:

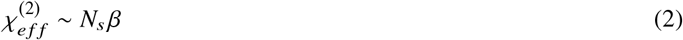

where *N*_*s*_ is the number of molecules contributing to the SHG process and *β* is the second-order hyperpolarizability. A significant increase in SHG signal is observed when electronic states in the molecules being probed are resonant with either the fundamental or second harmonic field. The coherent nature of SHG, which gives rise to the constructive and destructive interference, also provides the unique capability to monitor the relative populations of molecules on the outer vs. inner leaflet of a lipid bilayer. Initial adsorption to the outer leaflet will cause an increase in SHG signal while molecules that have flipped into the inner leaflet will generate SHG with the opposite phase leading to destructive interference of the detected signal over time (Fig. 1a). In addition to changes in the population of molecules in different positions within the membrane, the sensitivity of SHG to *β* also yields information about the local environment. Changes in solvent and aggregation states can change the observed hyperpolarizability by reported factors of up to 6 (30–32), which can subsequently increase or decrease the detected SHG signal by over an order of magnitude (Fig. 1b).

**Figure 1:**
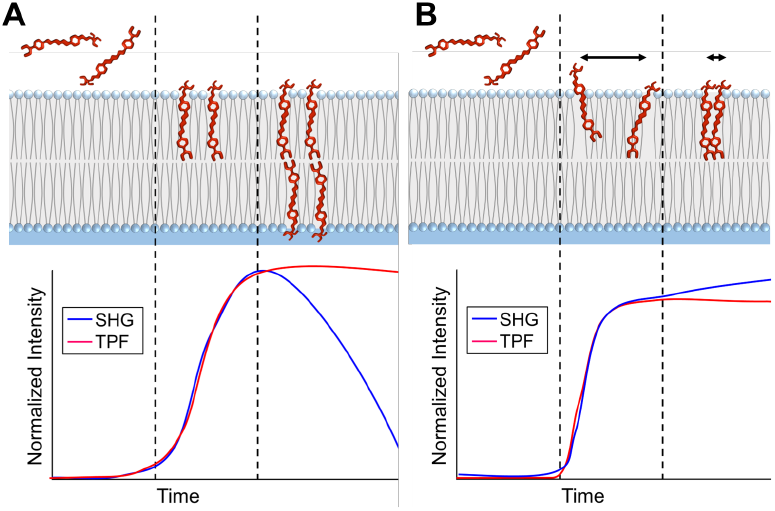
Schematic of SHG/TPF signals from FM 4-64 molecules interacting with a lipid bilayer over time. (A) SHG and TPF signals corresponding to probe molecules exhibiting flip-flop in the membrane -(i.) before probe molecules insert into membrane, (ii.) after probes insert into the outer membrane leaflet, and (iii.) when probe molecules flip into the inner leaflet and align with molecules on the outer leaflet. (B) SHG and TPF signals corresponding to probe aggregation on the outer leaflet of the membrane -(i.) before probe molecules insert into membrane, (ii.) after probe molecules insert into outer membrane leaflet, and (iii.) when probe molecule begin to cluster together on outer membrane leaflet over time.

For our experiments on *E. faecalis* and *S. aureus*, we chose the styryl membrane probes FM 4-64 and FM 2-10 (Fig. 2) to not only compare the different membranes, but also their sensitivity to relatively small molecular changes. These FM molecules contain identical dicationic headgroup regions with similar hydrophobic tails. FM 4-64 has been used for imaging and exhibits strong SHG signal when monitoring neuron membrane dynamics (33–37). FM 2-10 is only different from FM 4-64 in the shorter length of its conjugated region as shown in Figure 2.

**Figure 2:**
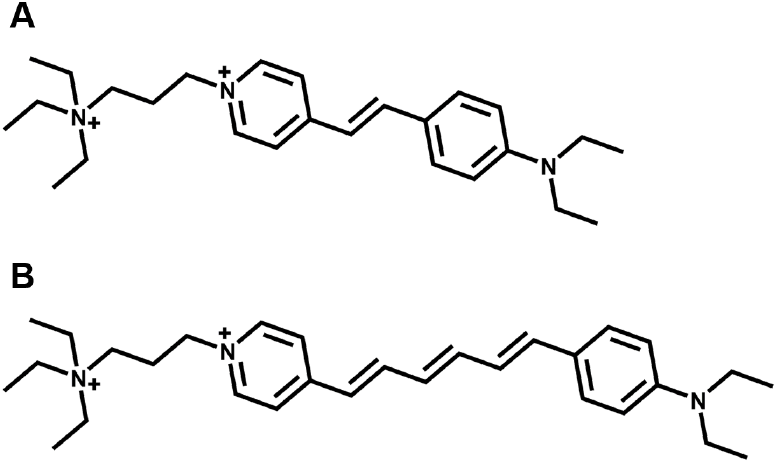
Structures of the membrane probe molecules used in the experiments. A. FM 2-10 and B. FM 4-64.

For each of these probe molecules, adsorption isotherms were collected with *E. faecalis* and *S. aureus* to assess the overall membrane affinities through the evaluation of adsorption equilibrium constants. In addition, the SHG signal was acquired over a two hour window after the probe molecules were introduced to the cell culture samples to monitor the dynamic changes. All of the studies were completed with living cells in early log phase (OD_600*nm*_ ~ 0.2) with rich media to minimize deviations from natural conditions. Our results show distinct behavior not only between the bacterial species, but also between the probe molecules despite their structural similarity. For both probe molecules, the SHG signal showed greater variations over time for *S. aureus*, but differences between how the signal changed is ascribed to different actions by the probes.

## MATERIALS AND METHODS

### Bacterial Strains and Probe Solutions

For these analyses, we used bacterial strains *Staphylococcus aureus* ATCC 27217 and *Enterococcus faecalis* OG1RF. Single colonies from brain heart infusion (BHI) agar plates were inoculated in 10 mL of BHI media and grown statically at 37°C overnight. The saturated cell cultures were then resuspended in fresh BHI media (Sigma-Aldrich, St. Louis, MO) to an optical density at 600 nm (OD_600_) of 0.01 and grown statically at 37° until reaching an OD_600_ ~ 0.20. The cells were then washed with sterile phosphorus buffered saline (PBS) via vacuum filtration and resuspended in BHI supplemented with 2% OxyRase™(OxyRase,Inc., Ontario, OH).

FM 4-64 (SynaptoRed™C2) and FM 2-10 (SynaptoGreen™C2) were purchased from Biotium (Fremont, CA) to use as membrane probes. Dye stock solutions were made with 80:20 sterile MilliQ deionized water (Millipore Sigma, Burlington, MA, 18.2 MΩ cm) to dimethyl sulfoxide (DMSO) (Fisher Scientific, Suwanee, GA) in order to minimize the final DMSO concentration to 0.2%.

### Flow Cell Apparatus

A home-built circulating gravity flow a pparatus w as i mplemented t o m inimize t he v olume o f s ample n eeded w hile also eliminating oscillation effects inherent to the peristaltic pump. The cell solution was pumped with a peristaltic pump from the sample reservoir into an elevated reservoir using biological grade tubing. From the elevated reservoir, the solution was allowed to freely flow through the laser focal region within a 2 mm path length quartz flow cell (Starna Cells Inc, Atascadero, CA) and recollect back into the initial sample reservoir. The dye was injected and mixed into the sample reservoir via a third port. Flow rate was 3 mL/s. The flow cell was prepped for pacification using a bovine serum albumin (Sigma-Aldrich, St. Louis, MO) and 0.1% glutaraldehyde (Fisher Scientific, Suwanee, GA) crosslinking protocol (38). The prepped cell cultures were added to the sample reservoir in the flow apparatus and baseline signals were collected for approximately 5 minutes before adding the FM probes to a final concentration of 16 *µ*M (with 0.2% final DMSO concentration).

### SHG/TPF Spectroscopy

Simultaneous SHG/TPF time-lapse signals were collected using a home-built spectroscopy instrument as shown in Fig. 3. The 80 MHz output of a MaiTai titanium:sapphire oscillator (Spectra Physics, Santa Clara, CA) centered at 800 nm was used to excite the sample. Pulse widths were compressed to ~78 fs and the average power at the sample position was adjusted to be 35 mW using a power attenuator consisting of a half-wave plate (*λ*/2 WP) and a polarizing beam splitting cube (PBSC) (ThorLabs, Newton,NJ). The laser was focused onto the sample using a 50 mm focal length lens (ThorLabs, Newton, NJ) paired with a 450 nm long pass filter to remove any SHG signal prior to the sample (Edmunds Optics, Barrington, NJ). A series of lenses (ThorLabs, Newton, NJ) were used to collect, collimate and focus the SHG and TPF signals onto a multimode fiber (Ocean Optics, Inc., Largo, FL). A 725 nm short pass filter was placed before the fiber to filter out the fundamental 800 nm from the SHG and TPF signals (Edmunds Optics, Barrington, NJ). The signals were then collimated upon exiting the fiber before being separated with a 450 nm longpass dichroic mirror (Edmunds Optics, Barrington, NJ). The SHG signal was isolated using a 400/10 nm bandpass filter and the TPF signals were filtered using either a 695/55 nm or 625/50 nm bandpass filter (Edmunds Optics, Barrington, NJ) for FM 4-64 and FM 2-10, respectively. The signals were simultaneously collected on photon counting photomultiplier tube (PMT) detectors (Hamamatsu, Bridgewater,NJ). The stability of the laser was monitored throughout the experiments by analyzing the SHG output from exciting a barium borate (BBO) crystal (Eksma Optics, Vilnius, Lithuania) with secondary laser line from the PBSC. The laser monitor signals were collected on a photodiode (PD) (ThorLabs, Newton, NJ). Data acquisition was controlled with home-built codes using LabVIEW (National Instruments, Austin, TX). Signals were collected for 8000 s at a sample integration time of 25 ms.

**Figure 3:**
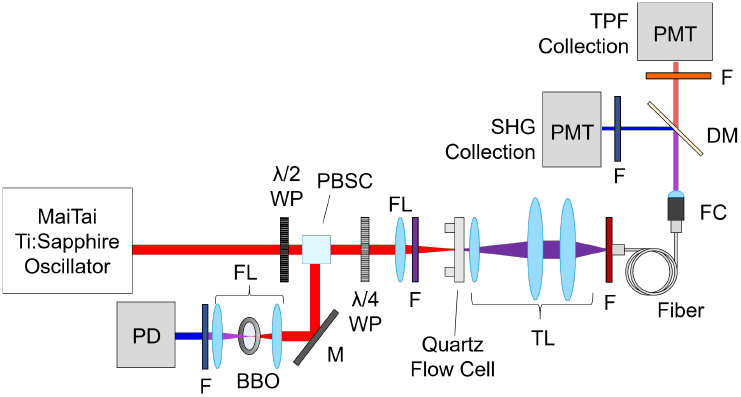
Schematic of SHG/TPF spectroscopy instrument. *λ*/2 WP: half-wave plate; PBSC: polarizing beam splitting cube; *λ*/4 WP: quarter-wave plate; FL: focusing lens; F: filter; TL: tube lens; FC: fiber collimator; DM: dichroic mirror; PMT: photomultiplier tube detector; M: mirror; BBO: barium borate cyrstal; PD: photodiode.

### Isotherms

Cell cultures, prepped as described above, were divided into 990 *µ*L aliquots. Dye stock solutions were made in various concentrations to keep the final concentration of DMSO to 0.2%. 10 *µ*L of dye was quickly mixed in a fresh aliquot before filling the flow cell with the sample. After collecting the SHG and TPF signals, the sample was removed from the flow cell in preparation for the next sample to be tested. Initial SHG/TPF signals were collected every 25 ms and averaged over 1 sec. The 0 *µ*M samples were used as solvent controls with 10 *µ*L of 80:20 sterile MilliPore DI H_2_O:DMSO added.

### Measurement of Cellular Viability

After the probe isotherms were conducted, 1:10 serial dilution growth plates were performed on BHI agar plates to test cell viability using the final probe concentrations of 0 *µ*M, 8 *µ*M and 16 *µ*M when the dye was first introduced to the cell sample (*t*_0*min*_), one hour (t_60*min*_), and two hours (t_120*min*_) after initial dye interaction. Since the spectroscopy experiments were conducted at room temperature, we tested for temperature variation effects by incubating half of each concentration sample at 37°C and half at room temperature for the time points t_60*min*_and t_120*min*_.

### Monitoring Bacterial Growth

To test temperature effects on the cells in the absence of FM probes, growth curves at OD_600_ were performed using a UV-Vis spectrometer (SHIMADZU UV-2600, Shimadzu, Columbia, MD) with 700 *µ*L cuvettes (Chemglass, Vineland, NJ). Overnight cultures are diluted into 50 mL of fresh BHI medium to an OD_600*nm*_of 0.01 and grown statically at 37°C. When the cultures reached an OD_600_ of 0.2, cells were washed via vacuum filtration in sterile PBS and resuspended into fresh BHI with 2% Oxyrase to an OD_600*nm*_= 0.2. The cell cultures were split into two 10 mL samples and grown statically over a two hour period where one sample was grown at 37°C and the other sample was grown at 20°C. Cell densities were measured at 600 nm every 30 minutes for a two hour period.

### Cytochrome *c* Assays

The determination of cellular charge was via the method by Peschel *et al* (39). Briefly, overnight cultures of *S. aureus* ATCC 27217 and *E. faecalis* were diluted into 200 ml of fresh BHI medium to an OD_600*nm*_of 0.01 and grown statically at 37°C. When the cultures reached an OD_600_ ≈ 0.3, cells were harvested via centrifugation, washed with 20 mM MOPS buffer, and concentrated to be an OD_600_ of 7 in 20 mM MOPS buffer. Cytochrome *c* (0.75 mg, Sigma-Aldrich, St. Louis, MO) was added and the cells were incubated at room temperature for 10 min. The mixture was centrifuged and the supernatant read at an OD_530*nm*_to determine unbound cytochrome *c*. We noted that *E. faecalis* bound far more cytochrome *c* (less free cytochrome *c* in assay measurements) than *S. aureus* using the originally published assay conditions (39), consequently, we increased the amount used in this assay to 0.75 mg. Shown are the averages ± standard deviations for *n*=3.

## RESULTS

### Isotherms

To assess the adsorption affinities of the probes for each of the bacterial species, isotherms were collected with both the SHG (Fig. 4) and TPF (Fig. S1) data. In these experiments the initial SHG signal is recorded for a series of increasing concentrations of added probe. The dissociation constants, *K*_*d*_, presented in Table 1 were extracted by fitting the SHG isotherms to the Langmuir adsorption model:

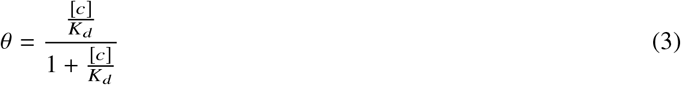

where *θ* is the coverage of probe molecules adsorbed to available sites on the membrane and [*c*] is the concentration of probe molecules in solution. The *K*_*d*_ values reflect the concentration of probe molecule at which half of the available sites on the membrane are occupied. As *K*_*d*_ is the equilibrium constant for dissociation, a higher numerical value denotes a weaker association between the molecule and the membrane.

**Figure 4:**
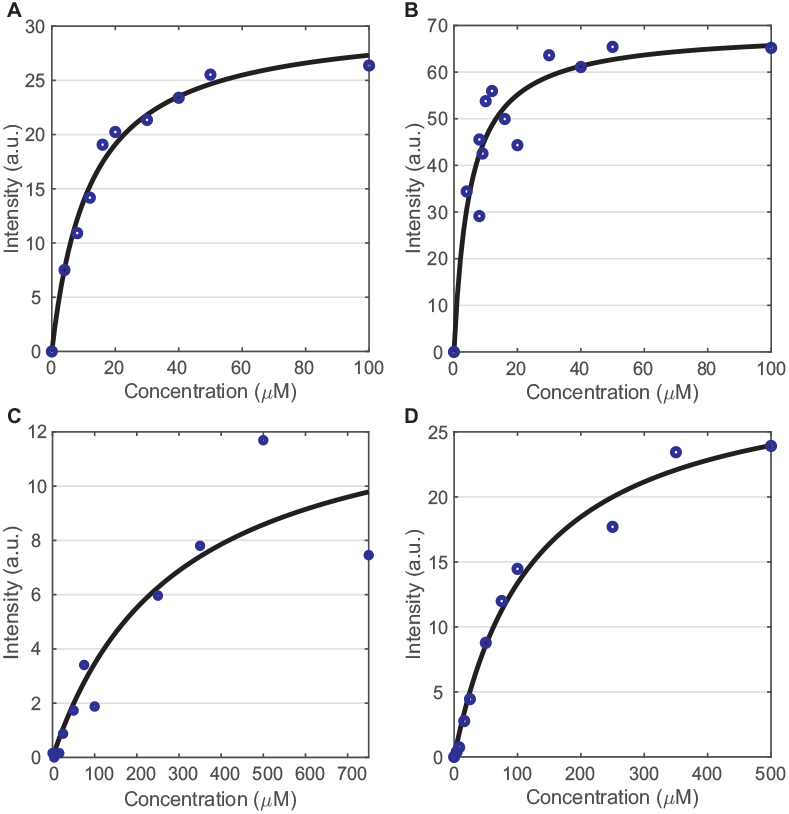
FM probe SHG isotherms. FM 4-64 isotherms for *S. aureus* (a) and *E. faecalis* (b). FM 2-10 isotherms for *S. aureus* (c) and *E. faecalis* (d). The dissociation constant, *K*_*d*_, is inset for each isotherm.

**Table 1:**
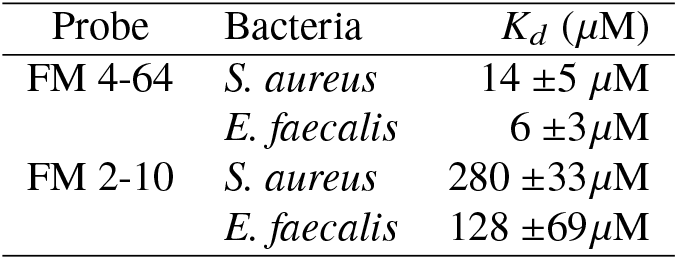
Average probe-membrane affinities in *S. aureus* and *E. faecalis*.

The data presented in Figures 4 and S1 represent the first time that adsorption isotherms of these molecules have been collected on *S. aureus* and *E. faecalis* using SHG and TPF, respectively. While adsorption isotherms on these systems have previously been reported from one-photon fluorescence measurements (40, 41), we also collected fluorescence-based isotherms by simultaneously monitoring the TPF (Fig. S1). The SHG measurements provide a more accurate reading of the molecules interacting with the cellular membranes. The noticeable differences and improved accuracy of the SHG relative to the TPF in our experiments is largely due to the increased two-photon fluorescence seen for the FM probes in media without cells present (Fig. S3). This increased fluorescence is likely due to interactions between the FM molecules and lipophilic clusters present in the media, as opposed to the phospholipids in the bacterial cell membranes, and therefore the fluorescence isotherms are also reporting on this affinity. SHG in contrast has a directional component to the signal emission that is dependent on the size of the scattering sample (11, 16, 42, 43). By detecting SHG in the forward direction in our experiments, we are preferentially collecting signal from sources larger than 500 nm, specifically the ~ 1 *µ* m b acterial c ells t hemselves, a s o pposed t o signal arising from smaller scattering bodies in the media which will predominately emit in the 90° direction (43).

Despite differences in the exact *K*_*d*_ values between our measurements and those reported from fluorescence measurements in the literature (41, 44), our measurements agree in that FM 4-64 generates smaller *K*_*d*_ values than FM 2-10 for both *S. aureus* and *E. faecalis*. This higher membrane affinity for FM 4-64 is not surprising given that FM 4-64 has a longer conjugation length than FM 2-10 (Fig. 2), making it a more lipophilic molecule for stronger adsorption to the bacterial membrane. Comparing between bacterial species, the *E. faecalis* membrane shows higher affinity for both probes. As discussed below, this may arise from the different surface charge on the membrane due to the composition of the phospholipid headgroups present.

### FM 4-64

To determine how the FM probe structures affect the bacterial membrane dynamics over time, SHG and TPF signals were simultaneously monitored over a 2 hour period. Three trials were conducted for each experiment with the average signal for the FM 4-64 shown in Figure 5 where the standard deviation is given by the highlighted regions. Looking first at the TPF in Fig. 5(b,d), both bacterial species generate similar signal profiles. There is first a large increase in signal as the dye is introduced into the system and adsorbs onto the membrane. The rate of the initial adsorption is not captured in these experiments as it occurs before the sample flows into the cuvette to be probed by the laser. As such, the rate of initial rise in the data is dictated by the flow rate of the sample solution through the system. After this initial rise, the TPF signal is relatively flat with a slight decrease over the 2 hour duration of the experiment. In order to confirm that the decrease in TPF signal for the FM 4-64 experiments (Fig. 5b,d) is due to photobleaching in our sample, a similar experiment was performed with FM 4-64 in *S. aureus* in which the incident 800 nm power was the same, but the flow rate of the sample was increased. As can be seen in Fig. S2, when the flow rate is increased, the TPF signal shows almost no decrease over the 2 hour duration of the experiment while the shape of the SHG signal remains unchanged, thus indicating the changes seen in the TPF signal is due to photobleach effects and not cell death.

**Figure 5:**
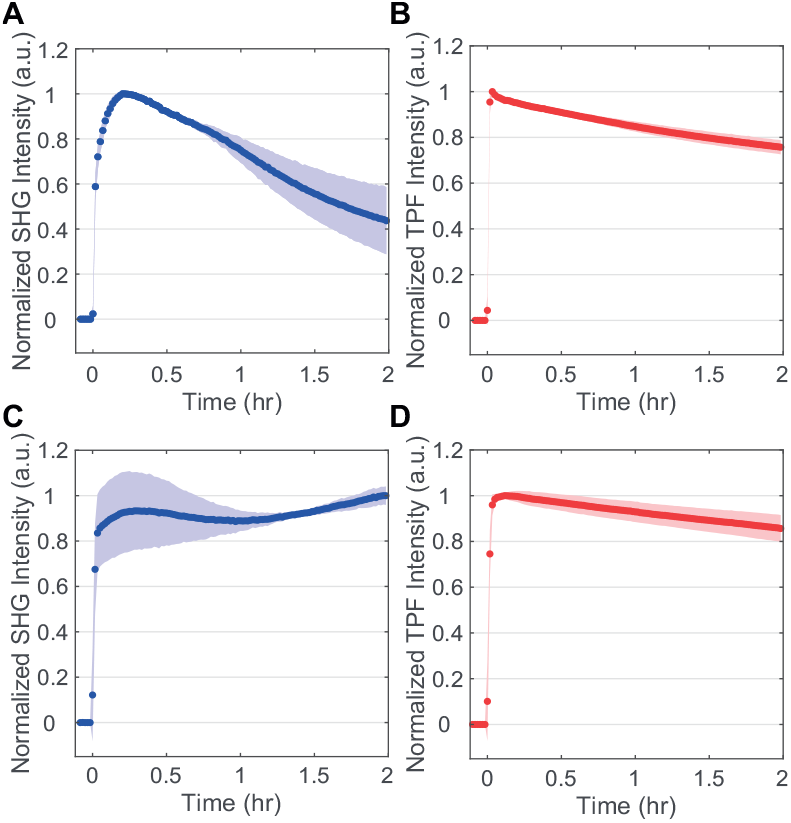
FM 4-64 SHG (a,c) and TPF (b,d) spectra while interacting with *S. aureus* (a,b) and *E. faecalis* (c,d) membranes. 16 *µ*M final concentration of FM 4-64. Standard deviations are shown as the shaded regions. *n*=3 for each plot.

In contrast to the TPF, the SHG signals in Fig. 5a,c show significant differences between the bacterial species. Over a two hour period, the SHG from FM 4-64 interacting with *E. faecalis* membranes (Fig. 5c) is static within the error of our measurement. In contrast, for *S. aureus* membranes, SHG signal from FM 4-64 decreases to approximately half of its initial intensity over the same time period.

In order to rationalize the time-dependent SHG signal, we consider multiple possible contributions to 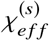 (Eq. 2) as the intensity of the incident light is unchanged during the time course of our experiments. One possible source of SHG decrease we ruled out is cell death. Given the relatively flat response of the fluorescence signal, we do not suspect appreciable changes in the population, *N*_*s*_. This is further supported by our assessment of cellular growth over the course of our experiments. Figure 6 shows the counts of colony forming units (CFUs) estimated from growth on agar plates of cells incubated with different concentrations of the FM molecules at different temperatures and times. The room temperature measurements with 16 *µ*M concentrations were extracted during the time course of the SHG measurements and therefore the cells were also subjected to the laser excitation and flow conditions. At these room temperature experimental conditions, *S. aureus* shows no change in the number of CFUs over the 2 hours of our SHG measurements. Similar results are shown for *E. faecalis* in the SI (Fig. S4). Further, these growth plate results show that the presence of the probe did not impact the ability to form viable CFUs. As such, we believe the stasis in our experimental conditions predominately arises from the 20°C temperature. Temperature-dependent growth measurements were also done by monitoring the OD_600_ of cells in the absence of probe molecules as shown in Figs. S5 and S6. Overall, given the high affinity of both probe molecules for the membranes and the lack of new membrane sites emerging, we conclude that changes in the number of probe molecules in the membrane are not the primary source of the changing SHG signal over time.

**Figure 6:**
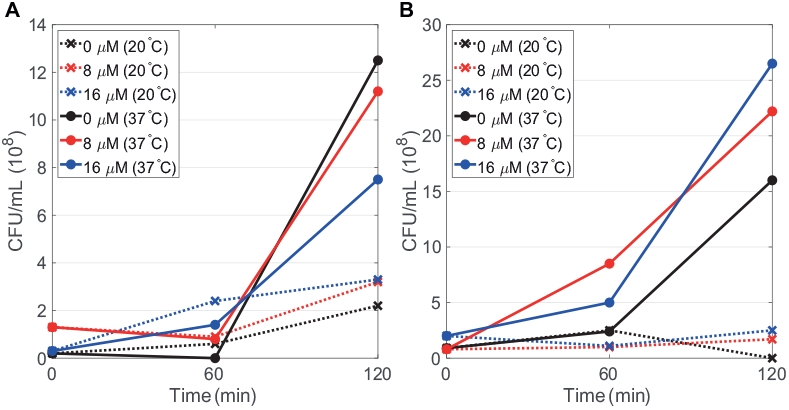
*S. aureus* viability effects when inoculated with FM 4-64 (A) and FM 2-10 (B) at 0 *µ*M (black), 8 *µ*M (red) and 16 *µ*M (blue) final probe concentrations when incubated at room temperature (crosses, dashed lines) versus 37°C (filled circles, solid lines).

Two other prominent mechanisms of SHG signal decrease that have been observed in living cells and do not require changing the number of molecules in the membrane are ion flux (11) and flip-flop (21, 28, 37, 45). A reduction in the SHG signal due to ion flux would arise from a loss of membrane potential; however, such a loss would also be expected to affect cell viability (46). Given that the growth plates do not show a reduction in CFUs during the course of the SHG experiments, we rule out ion flux as the source of the SHG signal decrease in *S. aureus*. This leaves flip-flop of the FM 4-64 in the *S. aureus* as the dominant mechanism leading to the dynamic SHG signal. As described earlier, the accumulation of probe molecules in the inner leaflet of the membrane will generate SHG signal with the opposite phase of that from molecules on the outer leaflet leading to destructive interference and an overall decrease in the measured SHG response over time. (Fig. 1a) The fact that flip-flop dominates in *S. aureus* but not *E. faecalis* also arises from significant differences in the membrane compositions of these bacteria as discussed below.

### FM 2-10

Turning to the time dependent interaction of FM 2-10, we see different behavior in all experiments despite the relatively small modification to the probe structure. In the TPF data shown in Fig. 7(b,d), we again see a fast initial rise dominated by the flow rate of our system, and the signal shows minimal decay over 2 hours. This probe molecule exhibits less photobleaching in our experiments relative to the FM 4-64 which is mostly likely due to a weaker two-photon cross-section at the 400 nm resonant wavelength (47).

**Figure 7:**
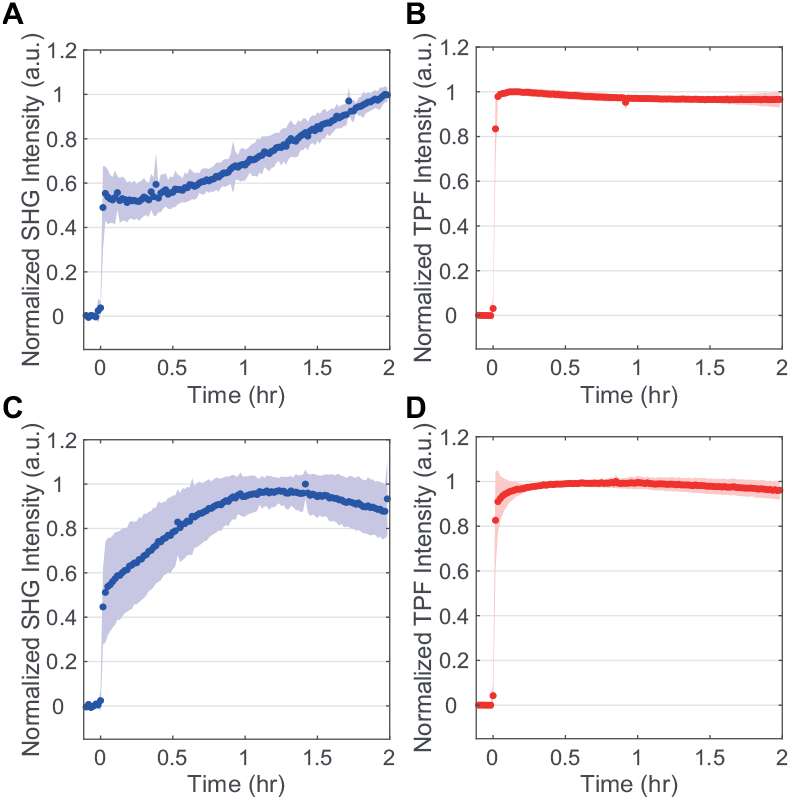
FM 2-10 SHG (a,c) and TPF (b,d) spectra while interacting with *S. aureus* (a,b) and *E. faecalis* (c,d) membranes. 16 *µ*M final concentration of FM 2-10. Standard deviations are shown as the shaded regions. *n*=3 for each plot.

The SHG dynamics for FM 2-10 (Fig. 7(a,c)) are again different from the TPF behavior, and more importantly, differ greatly from the FM 4-64 SHG data. After the initial rapid rise from membrane adsorption and flow, as lower rise is observed for both bacterial species. This slower increase in the signal levels off at approximately one hour for *E. faecalis* while in *S. aureus* the rise in SHG signal continues throughout the duration of the experiment. As discussed above, the rise cannot be due to an increased population of membrane bound species due to cell growth which also matches the static TPF signal. In this case then, the rise in SHG is attributed to the hyperpolarizability component of the signal (Eq. 2). As stated previously, large increases in the hyperpolarizability have been shown under different environmental conditions. Specifically, enhancements up to a factor of 6 have been observed for aggregation (30, 31) while the solvent environment alone can alter the observed hyperpolarizability by a factor of 2 (32). As detailed below, this behavior is indicative of either aggregation or translocation of the probes molecules to more ordered domains in the membrane.

## DISCUSSION

Two main contributing factors to how phospholipids regulate molecules permeating and transporting through the membrane are the headgroup charge and acyl tail structure. *S. aureus* and *E. faecalis* are known to have overall negatively charged membranes due to phospholipid headgroups such as phosphatidylglycerol (PG, −1 charge) and cardiolipin (CL, −2 charge). However, the overall membrane charge can fluctuate with the inclusion/substitution of positively charged phospholipids such as lysyl-PG (LPG, +1 charge) and neutral phospholipids such as digalactosyldiacylglycerol (DGDG, neutral). Such fluctuations are not only seen between bacterial species, but can change within a single species based on environmental stressors. For example, more cationic antibiotic resistant bacteria have been shown to change their overall membrane surface charge from negative to neutral, and in the most resistant strains, to positive (48). The membranes of *S. aureus* have been shown to be composed primarily of PG and LPG phospholipids while *E. faecalis* has been shown to contain PG, CL, DGDG and some LPG phospholipids (5, 49). In order to further support the difference in surface charge expected between the bacterial species, we examined whether there were overall differences in the abilities of *S. aureus* and *E. faecalis* to bind the positively charged (cationic) protein cytochrome *c* using a previously described method by Peschel *et al* (39). There was significantly more binding of cytochrome *c* to *E. faecalis* in comparison to *S. aureus* (Fig. S7). While these measurements are reflective of overall cell envelope charge (including cell wall and cell membrane), an overall higher ratio of negatively charged cell envelope in the *E. faecalis* membranes (5) is expected to be the source of the increased affinity for this species to the dicationic probes shown in the SHG isotherms (Fig. 4).

When considering the time-dependent response of the probes in our systems, the SHG signal from *S. aureus* showed greater change than *E. faecalis*, especially at longer time periods. Given that the cell samples are prepared at 37°C, but the experiments are performed at room temperature, ~ 20°C, the behavior at longer times likely derives from the cells’ response to the temperature change. These differences in temperature response for the two bacterial species may be attributed to differences in the fatty acid tails of the phospholipids which are integral to establishing the degree of membrane fluidity. More fluidic membranes are typically composed of shorter acyl chains (50), along with chains that are branched or unsaturated (51, 52). *S. aureus* is known to have a greater percentage of branched acyl chain structures in its lipidome in comparison to *E. faecalis* (4, 48, 49, 53, 54). In addition, the CL headgroup that is more prevalent in *E. faecalis* attaches four chains, as opposed to two chains for the other headgroups mentioned, which could also contribute to altered fluidity (5, 49, 55). While it is well known that bacteria do change their lipid tail composition in response to environmental temperature (2), the potentially more fluid membrane environment of *S. aureus* due to branched acyl chains may lead to the more dynamic behavior of the probes in these membranes over longer time periods.

For the specific experiment of FM 4-64 with *S. aureus*, we attribute the decrease of SHG signal at longer times to the flip-flop of the probe molecule from the outer to inner leaflet. The expected rate of flip-flop is not well known as a large number of different experiments on different systems has produced a wide range of results (21, 28, 56–64). It should be first noted that FM probes have been reported to be permanently bound to the outer leaflet of a bilayer membrane due to their double positive charge (65), incapable of translocating to the inner leaflet without pore or endocytosis mechanisms (66). This is in contrast, however, to other SHG measurements that have been able to detect membrane flip-flop dynamics with FM 4-64 in human embryonic kidney cells (37). Further, other dicationic styryl probes such as di-4-ANEPPDHQ have been observed to flip-flop in model vesicles using SHG (67). The fact that we observe signatures of FM 4-64 flip-flop in *S. aureus* but not *E. faecalis* may also be explained by the differences in their membrane fluidity and phosopholipid head group charge. As discussed above, the membranes of *S. aureus* have more phospholipids with branched acyl chains than *E. faecalis*, and this has previously been shown to increase the membrane fluidity (51, 52). Previous theoretical and experimental studies have both shown that increasing the fluidity of the different membranes being investigated was directly correlated with an increase in the flip-flop rate observed. (68–70) Moving to the potential impact from differences in phospholipid head groups, it has been shown that electrostatic interaction between the translocating molecule and the charge of neighboring lipid molecules can affect the flip-flop rate (61, 70). As such, we believe that flip-flop is not observed in our measurements on *E. faecalis* due to both the presence of a less fluid membrane and its more negatively charged surface.

The dynamic interaction of FM 2-10 with *S. aureus* leads to an increase in SHG signal resulting from an increase in the molecule’s hyperpolarizability. For our system, this behavior could arise from membrane organization such as the translocation of the probes to a lipid domain of significantly different physical properties, analogous to a change in solvent. This could be, for example, a local environment exhibiting a higher dielectric constant as has been implicated in the case of organic solvents (32) or it could be a more rigid environment to enforce improved ordering of the aligned probe molecules leading to stronger constructive interference of the SHG. Alternatively, over the course of these measurements the molecules could be self-assembling into aggregate structures within the membrane (Fig. 1b). While we would expect such an aggregation to be dependent on the concentration of the probe molecules, we were unable to collect sufficient signal from the FM 2-10 experiments at much lower concentrations for the low power necessary to prevent degradation of the cell samples. As such, we can not presently distinguish between these possible mechanisms, and include the likely possibility that our signal is combination of these behaviors. We do, however, conclude that the rise in SHG signal over the two hour time period in these experiments is a result of larger scale organization within the membrane and that this is the first time SHG has been shown to be sensitive to these dynamics in living cells.

## CONCLUSION

The application of SHG spectroscopy for studying the interaction of the membrane probes, FM 4-64 and FM 2-10, with living *S. aureus* and *E. faecalis* cells in rich media has yielded numerous findings. First, our SHG adsorption isotherms confirm the higher membrane affinity for FM 4-64 over FM 2-10 and also show stronger association of *E. faecalis* membranes to these molecules. Second, our time-dependent SHG measurements reveal dramatically different behavior for each combination of probe and bacteria. This highlights the sensitivity of each membrane to small structural changes in the molecules they encounter while also demonstrating how affected such encounters are to differences in the membrane composition. For *S. aureus*, the probe molecules continue to alter their behavior and the resulting SHG signal at longer time periods due to a more dynamic membrane environment relative to *E. faecalis*. An observed decrease in the SHG signal for the longer FM 4-64 molecule in *S. aureus* is attributed to the translocation of this molecule to the inner leaflet of the membrane while a slow rise in the measured SHG for FM 2-10 derives from the movement to a more organized structure. The interpretations of all of these findings are consistent with the known compositions of the membranes of the bacteria. Finally, in addition to the biological implications for these systems, our results show for the first time how SHG can be sensitive to molecule self-assembly in the membranes of living bacteria.

## AUTHOR CONTRIBUTIONS

LNM, EMF, and TRC contributed to the design of the experiments. WTB, JDW and EMF completed and analyzed the cytochrome *c* experiment. LNM carried out all of the other experiments, and the resulting data was analyzed by LNM and TRC. LNM, EMF, and TRC helped prepare the manuscript.

## ACKNOWLEDGMENTS

We would like to thank John Harp for guidance will proper handling and analysis of the bacterial cultures and serial dilution growth plating. This work was funded by NIH/NIAID grant R01AI116571.

