## Supplemental Figures S1-S7 for "Second Harmonic Generation Spectroscopy of Membrane Probe Dynamics in Gram-Positive Bacteria"

### SUPPLEMENTARY MATERIAL

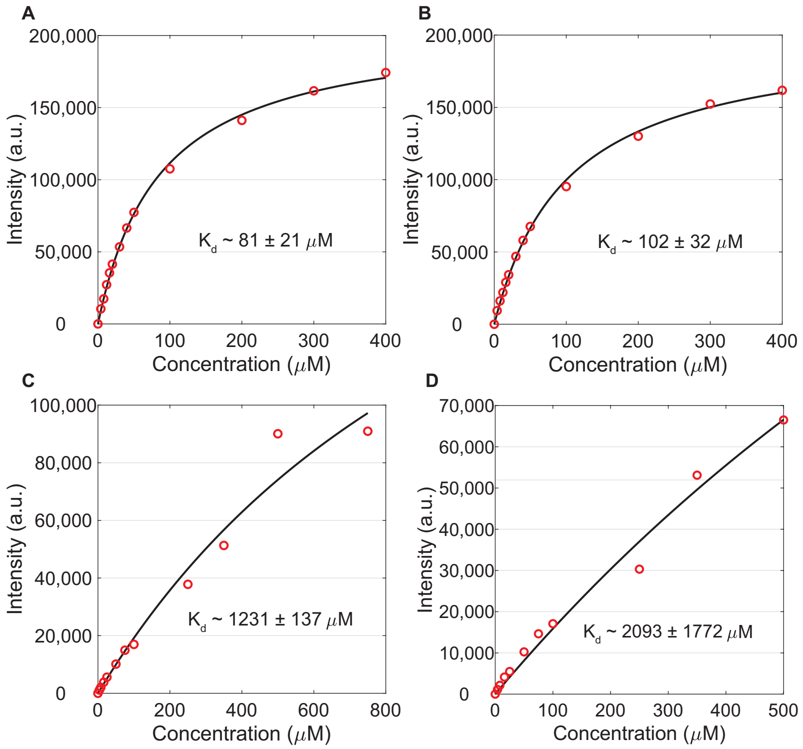


Figure S1: FM probe TPF isotherms. FM 4-64 isotherms for *S. aureus* (a) and *E. faecalis* (b). FM 2-10 isotherms for *S. aureus* (c) and *E. faecalis* (d). The dissociation constant, $K_d$, is inset for each isotherm.

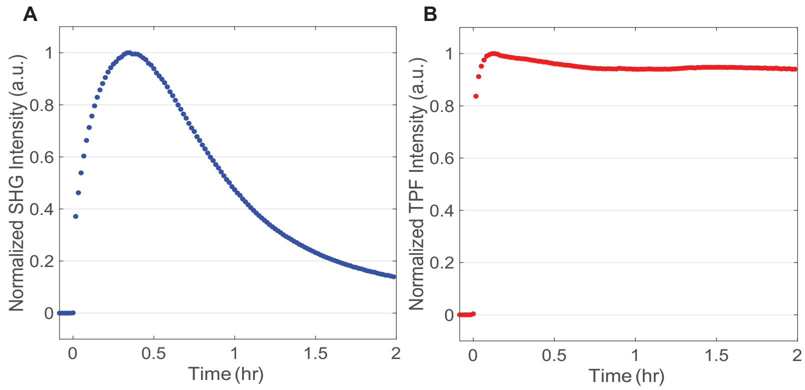


Figure S2: Faster flow rate effects on FM 4-64 SHG (A) and TPF (B) signals overtime in *S. aureus* cells.

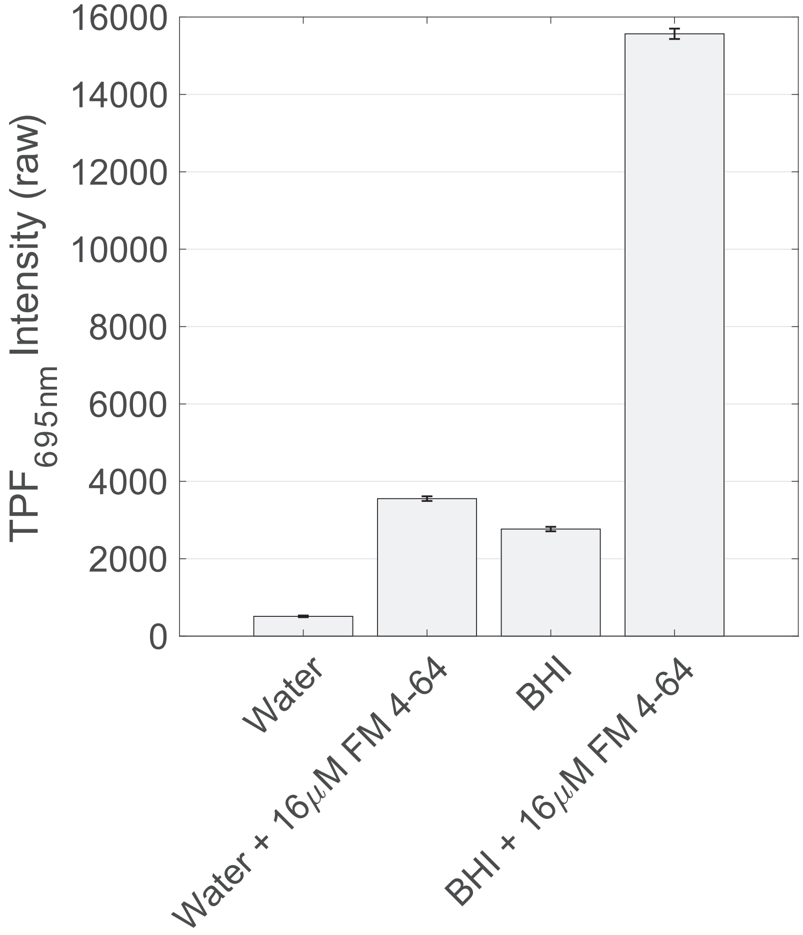


Figure S3: Background TPF comparison of water, water + 16 $\mu$M FM 4-64, BHI, BHI + 16 $\mu$M FM 4-64. TPF signal was collected using the instrumentation shown in Fig. 3 using the 695/55nm bandpass emission filter. The TPF signals were collected using a 25 ms integration time and averaged over 3 s.

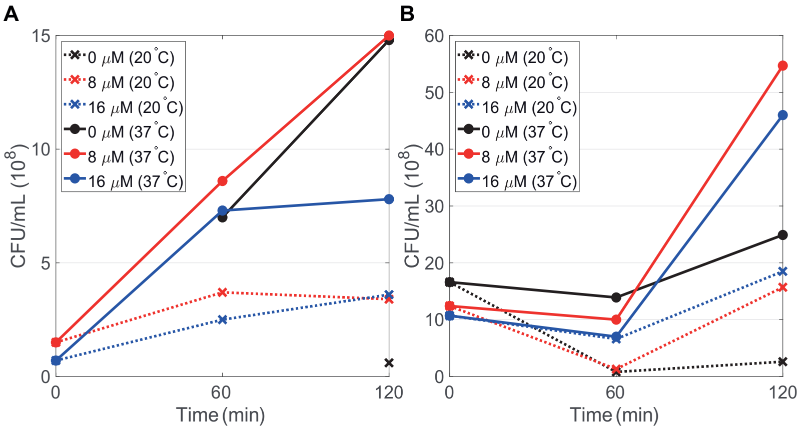


Figure S4: *E. faecalis* viability effects when inoculated with FM 4-64 (A) and FM 2-10 (B) at 0 $\mu$M (black), 8 $\mu$M (red) and 16 $\mu$M (blue) final probe concentrations when incubated at room temperature (crosses, dashed lines) versus 37°C (filled circles, solid lines).

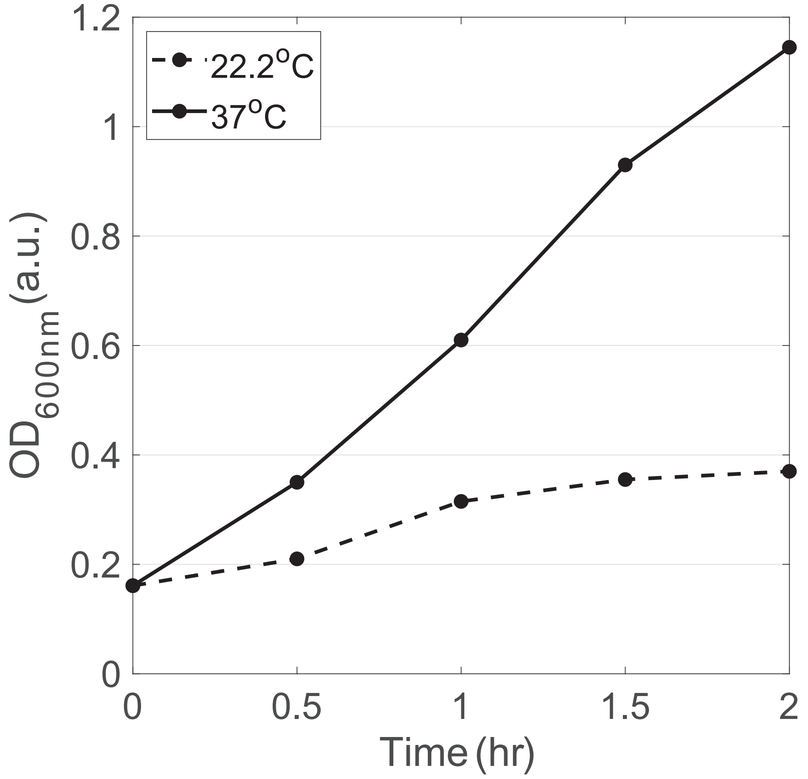


Figure S5: Temperature effects on growth rate of *S. aureus* cells starting at $t_{0min}$ with cell $OD_{600}$ ~ 0.2 for cells incubated at room temperature (dashed lines) versus 37°C over a two hour period.

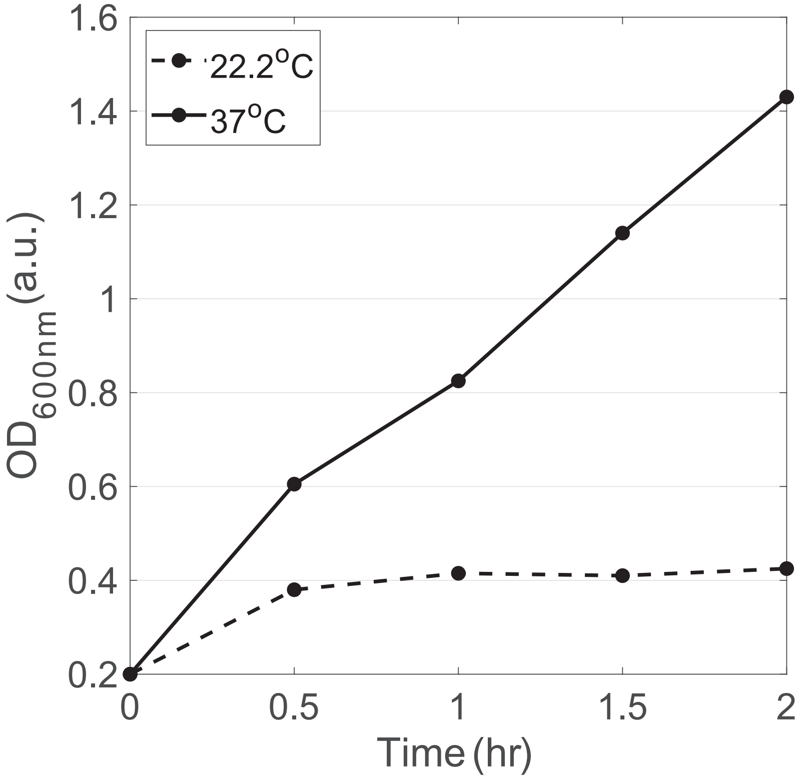


Figure S6: Temperature effects on growth rate of *E. faecalis* cells starting at $t_{0min}$ with cell $OD_{600}$ ~ 0.2 for cells incubated at room temperature (dashed lines) versus 37°C over a two hour period.

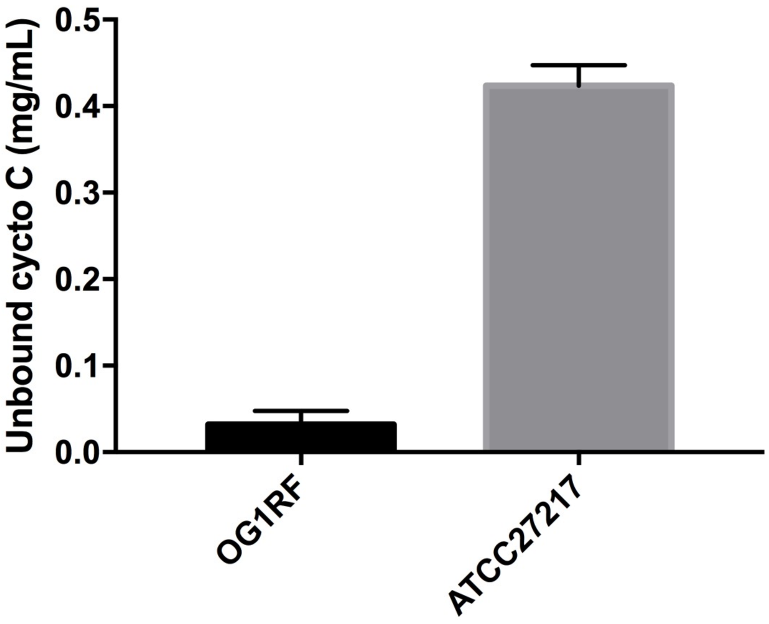


Figure S7: Interaction of cytochrome *c* to *E. facaelis* and *S. aureus* cells. Cells were incubated with equal amounts of the positively charged protein cytochrome *c*. Following centrifugation, the amount of protein within the supernatant (unbound) was determined at $OD_{530nm}$. Shown are the averages ± standard deviations for *n*=3.
